# Distinct chemotactic behavior in the original *Escherichia coli* K-12 depending on forward-and-backward swimming, not on run-tumble movements

**DOI:** 10.1101/2020.02.13.947150

**Authors:** Yoshiaki Kinosita, Tsubasa Ishida, Myu Yoshida, Rie Ito, Yusuke V. Morimoto, Kazuki Goto, Richard M. Berry, Takayuki Nishizaka, Yoshiyuki Sowa

**Affiliations:** Department of Physics, Gakushuin University, 1-5-1 Mejiro, Toshima-ku, Tokyo 171-8588, Japan; Department of Physics, University of Oxford, Park load OX1 3PU, Oxford, UK; Department of Frontier Bioscience and Research Center for Micro-Nano Technology, Hosei University, Tokyo 184-8584, Japan; Department of Physics and Information Technology, Faculty of Computer Science and Systems Engineering, Kyushu Institute of Technology, Iizuka, Fukuoka, Japan

## Abstract

Most motile bacteria are propelled by rigid, helical, flagellar filaments and display distinct swimming patterns to explore their favorable environments. *Escherichia coli* cells have a reversible rotary motor at the base of each filament. They exhibit a run-tumble swimming pattern, driven by switching of rotatory direction which causes polymorphic flagellar transformation. Here we report a novel swimming mode in *E. coli* ATCC10798, which is one of the original K-12 clones. High-speed tracking of single ATCC10798 cells showed forward and backward swimming with an average turning angle of 150°. The flagellar helicity remained right-handed with a 1.3 μm pitch and 0.14 μm helix radius, which is assumed to be a curly type, regardless of motor switching; the flagella of ATCC10798 did not show polymorphic transformation. The torque and rotational switching of the motor was almost identical to the *E. coli* W3110 strain, which is a derivative of K-12 and a wild-type for chemotaxis. The single point mutation of N87K in FliC, one of the filament subunits, is critical to the change in flagellar morphology and swimming pattern, and lack of flagellar polymorphism. *E. coli* cells expressing FliC(N87K) sensed ascending a chemotactic gradient in liquid but did not form rings on a semi-solid surface. Based on these findings, we propose a flagellar polymorphism-dependent migration mechanism in structured environments.

## Introduction

The flagellar motor is the most extensively investigated motility system in bacteria [1-3]. The motor complex is composed of approximately 30 different proteins and is attached to the helical flagellar filament via a hook structure. Most flagellar motors rotate in both directions, and the rotating filament works as a screw to generate thrust against the surrounding medium [4, 5]. Flagellated bacteria exhibit distinct chemotactic behaviors to move toward favorable environments. An *E. coli* cell has 5-10 left-handed flagellar filaments protruding from its cell body, and the rotation of a bundle of multiple flagella, rotating in the counterclockwise (CCW) direction (when viewed from filament to motor), propels a cell forward [4, 6]. The cell undergoes reorientation (tumbling) upon switching of flagellar rotation from CCW to clockwise (CW), which leads to a change in filament shape from left-to right-handed [7]. *V. alginolyticus* cells form a single left-handed polar flagellum, whose CCW and CW rotation propels a cell forward and backward, respectively [8, 9]. Additionally, cells of *V. alginolyticus* change their swimming direction by ∼90° due to a buckling instability of their straight hook (flick). Recently, a novel type of chemotactic behavior has been discovered, where the right-handed flagellum wraps around the cell body and propels the cell forward by its CW rotation [10, 11].

*E. coli* is an ideal model organism, due to rapid growth in pure nutrient media and abundant genetic strains. Theodor Escherich, the German pediatrician, isolated *Bacterium coli* from the feces of healthy individuals in 1885, which was renamed *Bacillus coli* and eventually *E. coli* in 1919 [12]. The original *E. coli* strain has been stored in the United Kingdom National Collection of Type Cultures as NCTC86, which shares a common genetic backbone with non-pathogenic *E. coli*, such as K-12, B and HS [12, 13]. In 1922, *E. coli* K-12 was isolated from the stool of a convalescent diphtheria patient [14]. Many hundreds of K-12 derivatives have been isolated for motility studies [15, 16], and K-12 has lost resistance to bacteriophage λ and sexual fertility (F^+^), from the effect of UV irradiation and acridine orange, during this period [14]. Initial studies isolated motile strains, such as W2637 and MG1655, using a semi-solid agar plate, in which motile strains formed a ring, but non-motile strains did not [17]. A recent study proposed a mechanism for navigated range expansion in *E. coli* cells, in a structured environment (semi-solid agar plate). It suggested a population fitness mechanism that recognizes nutrients and chemical gradients, which serve as a local guide and allow rapid expansion into unoccupied territories (outer edge) [18]. However, the reason why the original K-12 strain exhibits no motility remains unclear, and we infer that these non-motile cells have uncharacterized and exciting features to contribute to the field of bacterial flagellar studies.

To investigate further, we checked the swimming motility of the original K-12 strain, ATCC10798. It did not show a swarm ring on a semi-solid agar plate but could swim, in liquid medium, with a forward and backward movement like *V. alginolyticus*. The FliC(N87K) substitution seems to have prevented flagellar polymorphism and a consequent change of chemotactic behavior from a run-tumble to forward-backward movements. We found that ATCC10798 cells could not swim in structured environments, although they could swim toward the attractant in liquids. On the other hand, *E. coli* cells which showed run and tumble strategy freely moved with 180°-reversals to escape, if their route was blocked. From these results, we argue the importance of flagellar polymorphism for migration in structured environments.

## Material and Methods

### Bacterial strains

*E. coli* K-12 strain, ATCC10798 and W3110 were used in this study. Other mutants were listed in Supplementary Table 1. Cells were grown at 37 °C on 1.5 % (wt/vol) agar plate (010-08725; Wako) containing T-broth (1 % (wt/vol) Tryptone [Difco], 0.5% NaCl), and a single colony was isolated and resuspended to 10 ml volume of T-broth or LB (1 % (wt/vol) Tryptone [Difco], 0.5 % yeast extract [Difco], 0.5 % NaCl) liquid medium [7]. The cells were grown to an optical density of 0.4-0.7 at 600 nm with shaking at 30 °C (Supplementary Fig. 1).

### Construction of the *fliC* mutants

Plasmids and primers used in this study were listed in Supplementary Table 2. We purified the genomic DNA of ATCC10798 and amplified the *fliC* gene by PCR. The sequence difference of the *fliC* gene between ATCC10798 and W3110 strains was only at 87 residues.

To check the effect of the point mutation of 87 residues on the flagellar morphology, we performed two independent experiments: (i) the complementation of Δ*fliC* strain with the plasmid encoding FliC(N87K); (ii) the replacement of chromosomal *fliC* gene of ATCC10798 strain with the *E. coli* wild-type FliC. The plasmid encoding FliC(N87K) was constructed based on pYS10 encoding wild-type FliC. The mutation in *fliC* was generated by the “QuikChange” site-directed mutagenesis method using 1217_fliC(N87K)-f(QC) and 1218_fliC(N87K)-r(QC) listed in Supplementary table 2. The mutation was confirmed by DNA sequence analysis.

The strain was constructed using a λ Red recombination system with plasmid pKD46 encoding the Red system [19] and positive selection for the loss of tetracycline resistance [20]. The selectable tetracycline-resistance gene *tetRA* was amplified by PCR using primers of 0196_fliC-tetRA-F and 0197_fliC-tetRA-R listed in Supplementary Table 2. The *tetRA* cassette was replaced in the chromosomal *fliC* locus of the ATCC10798. After selection and isolation, SHU101 [*fliC*(N87K)::*tetRA*] was obtained and confirmed by colony PCR using 0219_fliC-(−175)-F, 0220_fliC-(+250)-R, 0210_tetRA-785-R and 0211_tetRA-1090-F (Supplementary Fig. 2). *tetRA* of SHU101 was replaced by the wild-type *fliC* of chemotactic wild-type strain RP437 amplified by PCR using primers of 1232_fliC-F and 0199_fliC-R. Tetracycline-sensitive clones were selected using tetracycline-sensitive plate and isolated as SHU102 [*fliC*(N87K)::*fliC*]. The strain construction was confirmed by sequence analysis using 0219_fliC-(−175)-F, 0220_fliC- (+250)-R, 0198_fliC-F and 0199_fliC-R.

**Figure 1.**
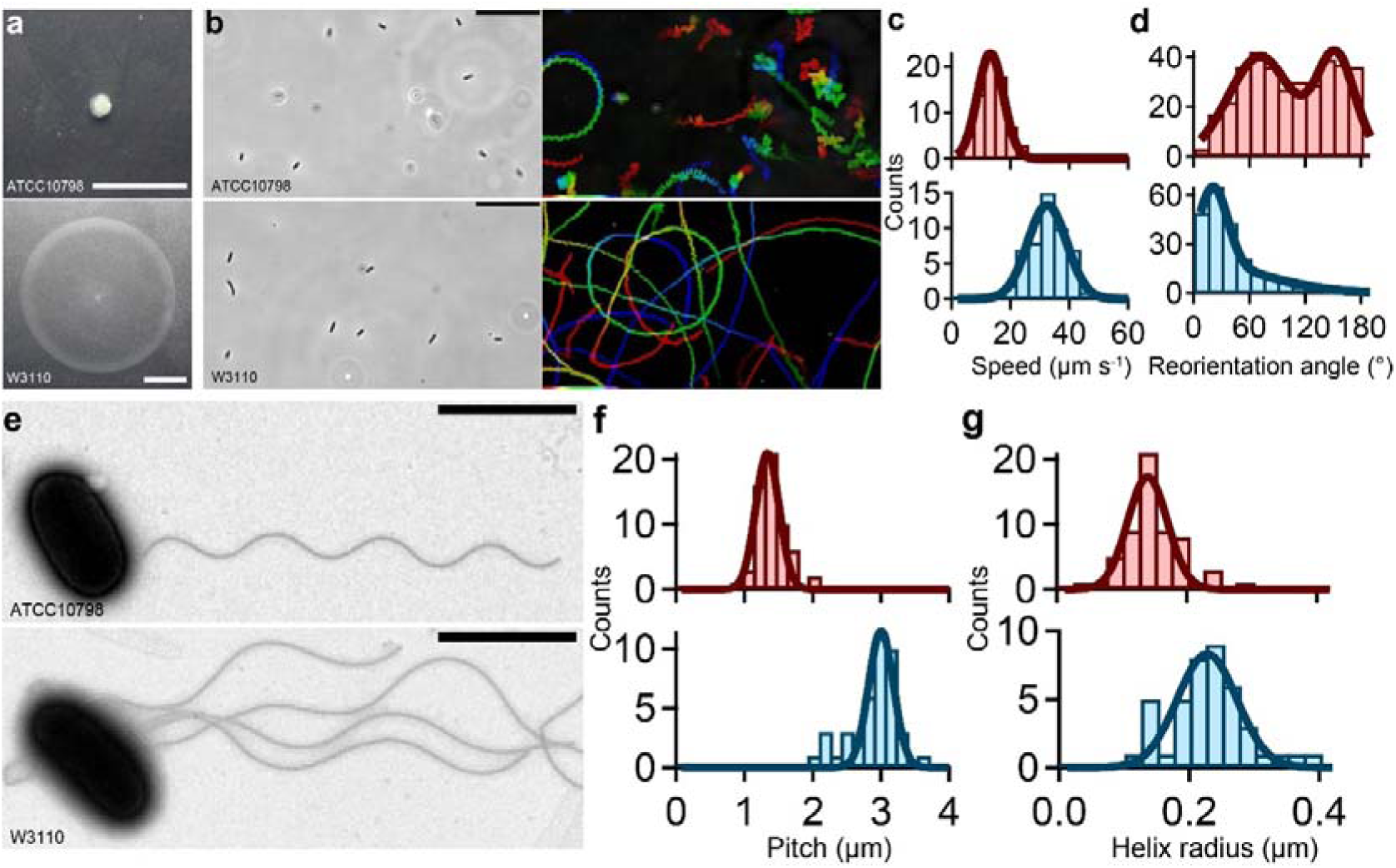
Characterization of swimming motility and structural parameters of ATCC10798 and W3110. (a) Motilities of *E. coli* ATCC10798 and *E. coli* W3110 cells on the 0.25 % (wt/vol) soft-agar plates at 30 °C for 7 h. Scale bar, 1 cm. (b) *Left*: Phase-contrast images. Scale bar, 20 μm. *Right*: Sequential phase-contrast images taken at 50-ms intervals throughout 10 s were integrated with the intermittent color code “red → yellow → green → cyan → blue.” (c) Histograms of the swimming speed of ATCC 10798 (*top*) and W3110 (*bottom*). The solid green lines represent the Gaussian fitting, where the peaks and SDs are 13.2 ± 4.4 μm s^−1^ in ATCC10798 (n = 70) and 32.5 ± 6.6 μm s^−1^ in W3110 (n = 50). (d) Histogram of the reorientation angles. The peaks and SD were 70 ± 31 degrees and 151 ± 23 degrees in ATCC10798 (n = 354) and 34 ± 13 degrees in W3110 (n = 119). (e) Electron micrographs of *E. coli* cells. Scale bars, 2 μm. (f) Histograms of the pitch. The solid lines represent the Gaussian fitting, where the peaks and SDs are 1.3 ± 0.2 μm in ATCC10798 (n = 59) and 3.0 ± 0.2 μm in W3110 (n = 41). (g) Histograms of the helix radius. The peaks and SDs are 0.14 ± 0.03 μm in ATCC10798 (n = 59) and 0.23 ± 0.05 μm in W3110 (n = 42).

**Figure 2.**
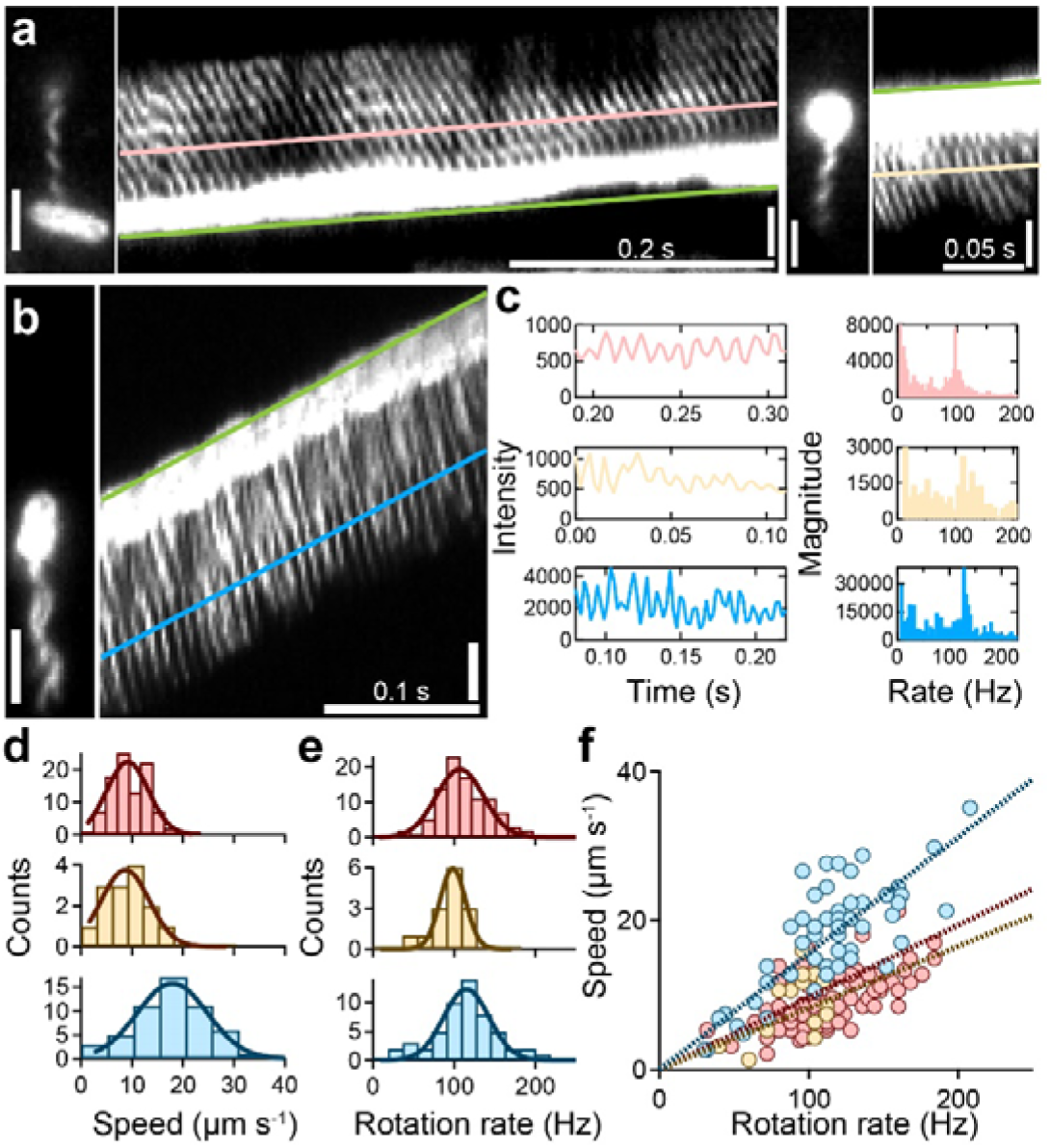
Visualization of forward and backward movements in ATCC10798. (a) Micrographs and kymographs of ATCC10798 cells during backward swimming (*left*) and forward swimming (*right*). The green line drawn at the tip of the cell enabled quantification of the swimming speed of the cell. The pink and blue lines were drawn on the signal of flagella, where the slopes were the same as that of a green line. Each intensity change indicated in Fig. 2c *left*. Scale bar, 2 μm. (b) Typical example of a run in W3110. Scale bar, 2 μm. (c) *Left*: The intensity changes along the flagellar filaments, whose each color corresponds to Fig. 2a and b. *Right:* The frequency analysis by a Fourier transform. The peaks in backward, forward swimming of ATCC1078 and run of W3110 were 94, 108 and 124 Hz, respectively. (d) Histograms of swimming speed of backward (*top*), forward (*middle*) in ATCC10798, and run in W3110 (*bottom*). The solid lines represent the Gaussian fitting, where the peaks and SDs are 9.1 ± 4.1 μm s^−1^ during backward swimming (n = 96), 8.7 ± 4.7 μm s^−1^ during forward swimming (n = 14), and 17.9 ± 6.9 μm s^−1^ during run (n = 54). (e) Histograms of the flagellar rotation rate. The peaks and SDs are 107.2 ± 29.1 Hz during backward swimming, 98.4 ± 15.7 Hz during forward swimming, and 115.1 ± 28.1 Hz during run. (f) Relationship between swimming speed and rotation rate. Each color corresponds to Fig. 2d and e. Dashed lines represent a linear fitting, with slopes of 0.083 μm per revolution during backward swimming, 0.097 μm per revolution during forward swimming, and 0.156 μm per revolution during run.

### Preparation of fluorescent-labeled cells

1ml of cultivated cells were collected by centrifugation at 6,000 ×g for 4 min at 25 °C, resuspended to buffer A (30 mM NaCl, 70 mM KCl, 2 mM EDTA) at pH 7.8 containing Biotin-NHS-ester (Dojindo), and incubated for 15 min at room temperature. After labeling, two rounds of centrifugation above mentioned removed excess biotin. Biotinylated cells were resuspended into buffer (30 mM NaCl, 70 mM KCl, 5 mM MgCl_2_) at pH 7.0 containing 0.1 mg/ml Cy3-conjugated streptavidin and incubated for 3 min [21, 22]. Two rounds of centrifugation removed excess dyes, and then cells were resuspended into buffer B.

### Electron microscopy

Carbon-coated electron microscope grids were glow-discharged with a hydrophilic treatment device (PIB-10; Vacuum Device) [10, 22]. Cells in buffer B were placed on the grid and incubated for 10 min at room temperature. Cells were chemically fixed with 2 % (vol/vol) glutaraldehyde in buffer B for 15 min. Cells were washed three times with buffer B and subsequently treated by 2 % (wt/vol) ammonium molybdate for staining. Samples were observed under a TEM (JEM-1400; JEOL) at 100 kV. Whole images were captured by a CCD camera as 8 bits.

### Motility assay on soft-agar plates

A single colony was inoculated on a semi-solid agar plate (0.25 % (wt/vol) T-broth soft-agar plates (214010; Difco)) and was incubated at 30°C for 7 hours. Ability of cell motility was evaluated from a colony’s diameter by Image J 1.45s (http://rsb.info.nih.gov/ij/).

### Motility assay

All experiments were performed at room temperature. The flow chamber was composed of two coverslips (no. 1, 0.12–0.17 mm thickness, Matsunami Glass) with different sizes (18 × 18 and 24 × 36 mm) [23, 24]. The 24×36 mm cover glass was glow-discharged with a hydrophilic treatment device (PIB-10; Vacuum Device) to clean its surface. Two pieces of double-sided tape, cut to a length of ∼30 mm, were used as spacers between coverslips. Two tapes were fixed with a ∼5 mm interval, and the final volume was ∼7 μl, indicating that the thickness of double-sided tape was ∼90 μm. In swimming assay, buffer C (30 mM NaCl, 70 mM KCl, 5 mM MgCl_2_, 5 mg/ml bovine serum albumin (BSA) [Sigma Aldrich]) was infused into the flow chamber, and then with 10 μl of the cell-suspension medium.

For observation of stuck cells under a total internal reflection fluorescence microscopy (TIRFM), a glass was coated with poly-L-lysine (F8920; Sigma Aldrich). Cells in buffer D (30 mM NaCl, 70 mM KCl, 5 mM MgCl_2_, 1.5 mg/ml BSA) were infused into the chamber, and then a 20 μl volume of buffer D was infused to remove unbound cells.

A capillary assay was performed with a method by Niikata et al [25]. We used a 10-μl tip as capillary, which contains 5-μl buffer B with 1 % (wt/vol) agarose. The tip was inserted into a chamber for a chemotactic response assay (Supplementary Fig. 6). Buffer C was infused into a chamber to prevent cells adhering to the glass surface.

For a tethered-cell assay, cell suspension with an optical density of around 0.6-0.8 at 600 nm was sheared by passing it back and forth 35 times between 1-ml syringes equipped with two 26-gauge needles connected by a peace of tubing. Cells were collected by centrifugation at 6,000 ×g for 2 min at 25 °C, resuspended to buffer E (10 mM KPi, 85 mM NaCl, 0.1 mM EDTA). After two rounds of washing, cells were resuspended into buffer E. Cells were stuck on a glass surface via an anti-FliC antibody with a 1: 300 dilution (Fig. 3a), and unbound cells were washed by buffer F (10 mM KPi, 67 mM NaCl, 0.1 mM EDTA, 10 mM lactate). Spinning cells were captured using a CMOS camera at 60 frames s^−1^ for 10 sec thorough 40× objective, as previously described [5, 26]. Rotational motions of cell bodies were analyzed using custom software based upon LabVIEW (National Instruments). The CW bias was defined as 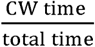 (Fig. 3c).

**Figure 3.**
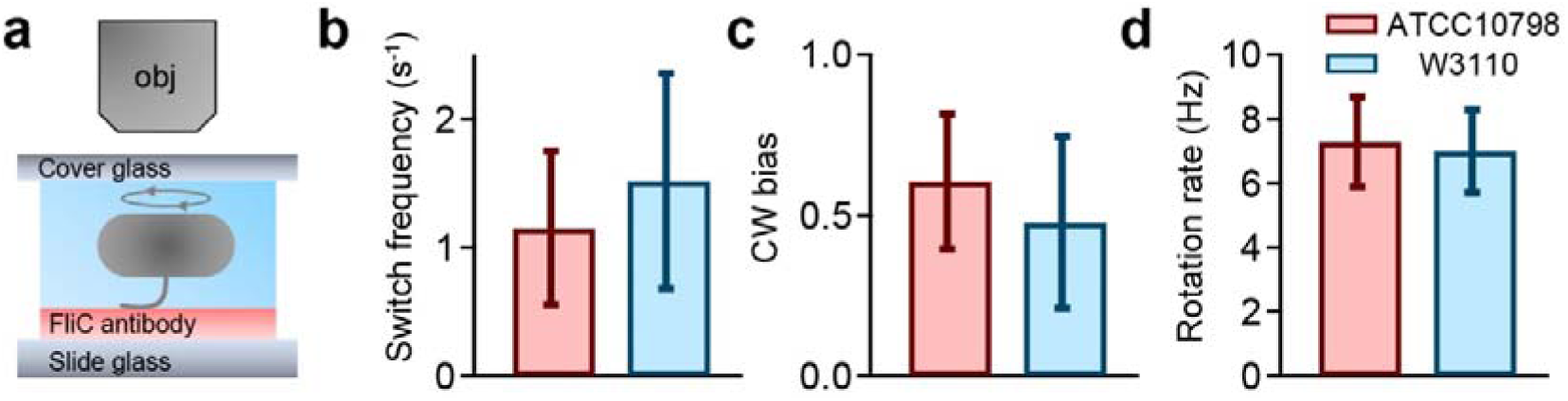
Quantification of switching behavior by tethered-cell assay. (a) Schematics of tethered-cell assay. Note that an upright microscopy is used in our measurement, meaning that the rotational direction of the motor is clockwise in the case of cell body rotating clockwise on the camera plane, vice versa. (b) Switching frequency of ATCC10798 and W3110 for 10 sec. The average and SD were 1.15 ± 0.60 s^−1^ in ATCC10798 (n = 99) and 1.52 ± 0.84 s^−1^ in W3110 (n = 53). (c) CW bias (Time_CW_/Time_Total_). The average and SD were 0.61 ± 0.21 in ATCC10798 (n = 99) and 0.48 ± 0.27 in W3110 (n = 53). (d) Rotation rates. The average and SD were 7.3 ± 1.4 Hz in ATCC10798 (n = 99) and 7.0 ± 1.3 Hz in W3110 (n = 53, *P* = 0.1864 > 0.05 by *t*-test).

### Microscopy

For visualization of fluorescent-labeled cells, a green laser beam (wavelength of 532 nm; Compass-315M-100, Coherent) was introduced into an inverted microscope (IX71, Olympus) equipped with a ×100 objective (Plan Apo TIRF, NA 1.49, Nikon Instruments), a dichroic mirror (custom-made, Chroma), an emission filter (NF01-532U, Semrock), an EMCCD camera (iXon+ DU860, Andor), a CCD camera (HR1540; Digimo), a highly stable customized stage (Chukousha) and an optical table (RS-2000, Newport). Images were recorded at 2.5-ms intervals, using an EMCCD camera with a magnification of 130 × 130 nm^2^ at the single pixel on the camera plate.

### Live-cell imaging on agarose

We conducted two independent experiments to investigate a swimming motility on a 0.2 % semi-solid agarose. First, we introduced 20 ml of 0.2 % agarose into a 4.4 cm radius of the Petri dish, inoculated a single-colony onto the agarose, and then incubated cells for 7 h at 30□ (Fig. 5a and b). Second, we introduced 20 μl of 0.2 % agarose onto a slide glass, wait until the agarose was solidified, and then put 10 μl of culture on it. The agarose pad was covered with a 22×22 coverslip using a double-sided tape with a ∼ 20 mm interval, and the approximate height is 75 μm, (Fig. 5c and d). W3110 cells near the bottom glass surface were observed to guarantee a swimming motility in agarose environments. Both experiments were carried using an upright microscope (Eclipse Ci; Nikon) equipped with a 40× objective (EC Plan-Neofluar 40 with Ph and 0.75 N.A.; Nikon), a CMOS camera (H1540; Digimo). Images were recorded at 20 fps for 15 sec.

### Data analysis

To identify reorientation events from trajectories, we used three strategies, as previously reported [27]. First, phase-contrast images were captured at up to 200 frames s^−1^. The centroid positions of cells determined swimming trajectories. Given the trajectory of cells, ***r* (*t*)** = [*x*(*t*), *y*(*t*)], the swimming velocity ***v*(*t*)** was defined as 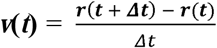 Second, to eliminate the effect of noise, such as Brownian motion, on reorientation events, we smoothed the data by calculating running averages over 10 points, which corresponded to 50-ms intervals. Finally, given the two data points, ***r* (*t*)** = [*x*(*t*), *y*(*t*)] and ***r*(*t +*** Δ***t*)** = [*x*(*t* + Δ*t*), *y*(*t* + Δ*t*)], we defined the angle against the horizontal axis as *θ* (*t*). If the two successive angle changes, *θ*(*t*_*1*_)-*θ*(*t*_*0*_) and *θ*(*t*_*2*_)-*θ*(*t*_*1*_) were over α, that point was identified as the end of the run. A new run begins at three successive angle changes < α with the speed of more 5 μm s^−1^; hence, the minimum duration of run was 80 ms. The threshold α is described as the following equation: α = c Δ*θ*_med_, where c the coefficient and Δ*θ*_med_ the median directional change. We manually checked the trace and video to avoid the detection of false events and found that the best value of c is 3.

Under fluorescent-labeled cell experiments, we constructed a kymograph at 2.5-ms intervals, as shown in Figures 2 to measure the swimming speed. The flagellar rotation rate of each cell was measured by Fourier transform analysis (Fig. 2c *right*). Under TIRF illumination, intensity changes were detected when fluorescent-labeled flagella made contact with an evanescent field. Intensity changes in a 2×2 pixel grid were measured and calculated by fast Fourier transform analysis [21, 22].

## Results

### Differences in swimming pattern and flagellar structure between *E. coli* ATCC10798 and W3110

We found that ATCC10798 cells did not form a swarm ring on the semi-solid agar plate, while W3110 cells were able to do so (Fig. 1a). Previous studies reported that cells defective in chemotaxis, motility or lacking active flagella did not form a ring on the semi-solid agar plate (Supplementary Fig. 2) [28]; therefore, it was conceivable that ATCC10798 cells would be unable to exhibit swimming or switching behavior. To address this, we performed microscopic measurement using a phase-contrast microscope (Supplementary Video 1). Unexpectedly, ATCC10798 cells showed swimming motility with reorientations, which is a different motility pattern to W3110 cells (Fig. 1b).

To better understand the different motility modes between strains, we quantified the swimming speed and switching pattern of cells. The average swimming speed ± standard deviation (SD) was 13.2 ± 4.4 μm s^−1^ in ATCC10798 cells and 32.5 ± 6.6 μm s^−1^ in W3110 cells (Fig. 1c). To characterize the switching behaviors in detail, we extracted angle changes between *θ*(*t*) and *θ*(*t* + Δ*t*) from an algorithm based on previous studies (see Material and Methods section) [27]. The frequency distribution of turning angle in ATCC10798 has a bimodal shape, with peaks at 70° and 150°. This indicates that cells reversed their swimming directions (Fig. 1d *top*), which was previously reported in *Salmonella enterica serovar*, a curly mutant of *S. typhimurium* [29]. The most frequent angles for changes of direction in W3110 cells were approximately 35°, which was similar to the angle previously reported (Fig. 1d *bottom*) [27].

Next, we analyzed flagellar morphology using TEM. In ATCC10798, the average number of filaments was two, and the average length ± SD was 4.7 ± 1.1 μm (Fig 1e *top*, n = 62). The flagellar pitch and helix radius were measured to be 1.3 ± 0.2 μm (Fig. 1f *top*) and 0.14 ± 0.03 μm (Fig. 1g *top*), respectively, which corresponded to the curly flagellar filament [30]. In W3110, cells formed approximately six flagellar filaments around the cell body (peritrichous flagella, Fig. 1e *bottom*). Their average length ± SD was 7.3 ± 1.9 μm (n = 48), and their pitch and helix radius were measured as 3.0 ± 0.2 μm (Fig. 1f *bottom*) and 0.23 ± 0.05 μm (Fig. 1g *bottom*), respectively. These helical parameters indicated that this flagellar filament belongs to the normal type [30]. Other structural parameters are summarized in Supplementary Table 3.

### Forward-backward movement in ATCC10798 swimming

To elucidate the basis for the difference in the swimming mode between ATCC10798 and W3110, we labeled the flagellar filaments with a fluorescent dye, Cy3, taking advantage of biotin-avidin interaction (see Material and Methods) [21, 22]. First, we observed the flagellar dynamics in a large field (230 μm × 144 μm). ATCC10798 cells, with few flagellar filaments, frequently exhibited forward and backward swimming, like *Vibrio alginolyticus* [8, 9], whereas cells with many filaments showed wobbling motion, apparently due to deficient bundle formation (Supplementary Video 2). Most W3110 cells exhibited directed linear motion (run) with abrupt directional changes (tumble) [6, 27].

We next observed the swimming speed and flagellar rotation rate simultaneously with a high S/N ratio at 400 frame s^−1^ (Supplementary Video 3) [21, 22]. Using a kymograph analysis (see Material and Methods), the swimming speeds and rotational rates were quantified from the slope and the changes in intensity, respectively (Fig. 2a-c). In ATCC10798, the swimming speed and rotation rate were estimated to be 9.1 ± 4.1 μm s^−1^ / 107.2 ± 29.1 Hz during backward swimming (Fig. 2a *left*, Fig. 2d and e *top*) and 8.7 ± 4.7 μm s^−1^ / 98.4 ± 15.7 Hz during forward swimming (Fig. 2a *right*, Fig. 2d and e *middle*). In W3110 (Fig. 2b), these parameters were 17.9 ± 6.9 μm s^−1^ and 115.1 ± 28.1 Hz (Fig. 2d and e *bottom*). Despite similar values of rotational rates between the two strains, flagellar rotation in W3110 cells was approximately twice as efficient: the ratio of swimming speed (*v*) / rotation rate (*f*) was 0.156 μm/rotation while that of ATCC10798 was 0.083 μm/rotation during backward swimming and 0.097 μm/rotation during forward swimming (Fig. 2f). This result suggests that the larger helix can produce a stronger thrust, as previously predicted by mathematical modelling [31].

### Real-time imaging of structure and kinematics for flagellar filaments, under TIRFM

We next determined flagellar structure and function, simultaneously, using TIRFM [10, 21, 22]. We found that a cell attached to the glass surface can rotate its flagellar filament freely, by treating coverslips with poly-L-lysine and BSA. In ATCC10798, we could see wave propagation, away from the cell body, during CW rotation of right-handed flagellar filaments, and towards the cell body during CCW rotation (Supplementary Video 4). From this analysis, we conclude that forward and backward movements in ATCC10798 cells are driven by CW and CCW rotation, respectively, for a right-handed flagellar filament. We summarized the flagellar morphology and rotation rate in both modes in Supplementary Fig. 3.

Although ATCC10798 cells had only right-handed flagellar filaments, W3110 cells had both right- and left-handed flagellar filaments (Supplementary Video 5). W3110 cells mainly formed left-handed flagellar helices when the filaments freely rotated in the CCW direction. The motor switching caused the gyration of the filament and transformation from the left-handed into right-handed filament within 100 ms (Supplementary Fig. 4 *left* and Supplementary Video 6). We also detected this reversible transformation from right-to left-handed (Supplementary Fig. 4 *right* and Supplementary Video 7). Furthermore, we observed coiled-state flagellar filaments with a radius of 0.78 ± 0.02 μm in W3110 (Supplementary Fig. 5).

**Figure 4.**
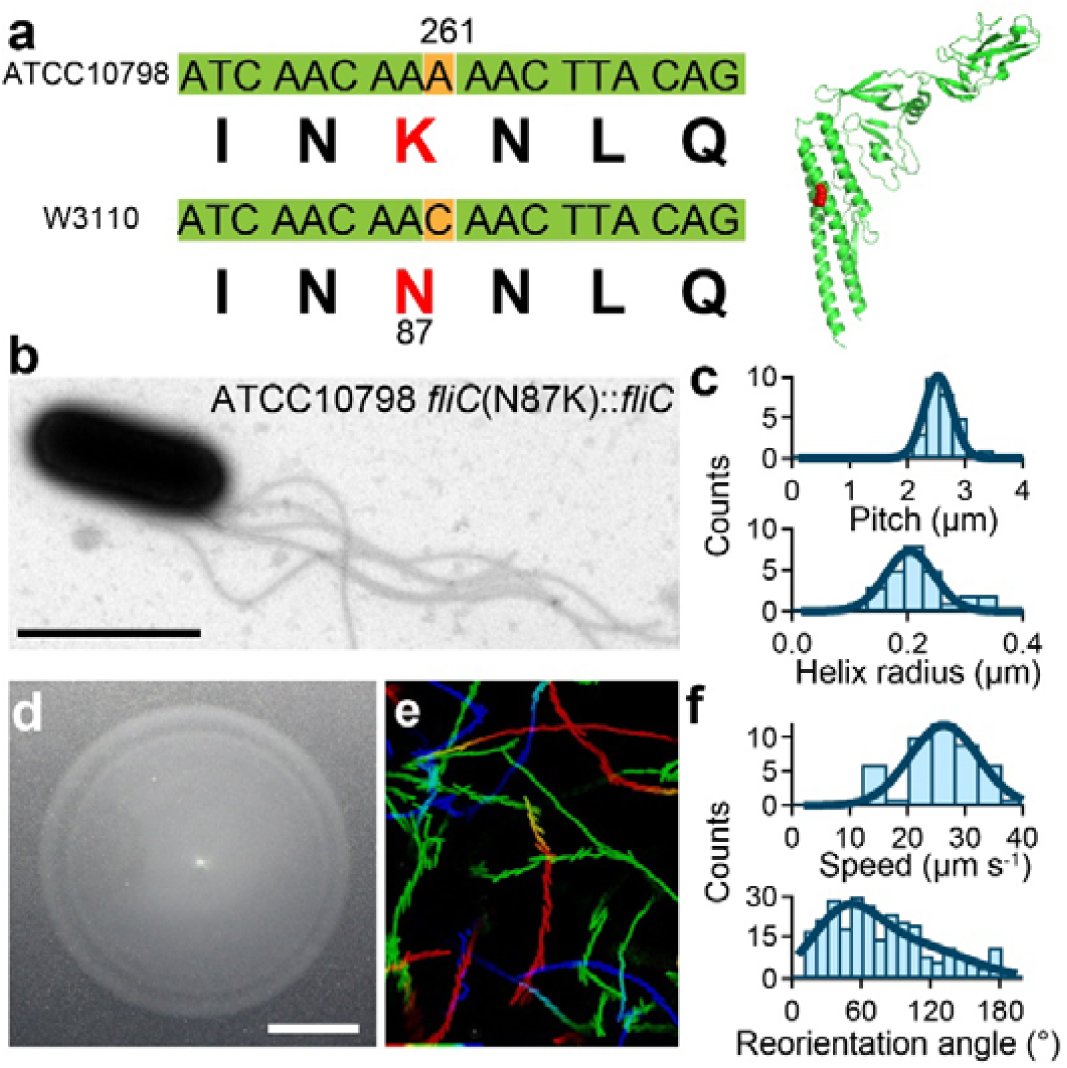
FliC(N87K) substitution alter flagellar shape and swimming mode. (a) *Left*: The gene sequence of the *fliC. Right*: The crystal structure of FliC (PDB: 1IO1). The red residue represents N87 molecule. (b) Electron micrograph of [*fliC*(N87K)::*fliC*] cells. Scale bars, 2 μm. (c) Histograms of the pitch (*top*, 2.5 ± 0.2 μm; n = 27) and the helix radius (*bottom*, 0.20 ± 0.04 μm, n = 27). (d) Motilities on a 0.25 % (wt/vol) soft-agar plates at 30°C for 7 h. Scale bar, 1 cm. (e) Swimming traces at 150-ms intervals for 15 s. The intermittent color code indicated the time course from red to blue. Area, 68.6 μm × 85.9 μm. (f) Histograms of the swimming speed (top, 26.3 ± 6.0 μm s^−1^, n = 45) and the reorientation angles (bottom, 46 ± 28 degrees, n = 269).

### Quantification of single motor behaviors by tethered-cell assay

Previous studies claimed to identify torque-dependent flagellar transformation, based on direct measurement using a dark-field microscopy and molecular simulation [32, 33]. However, we could not detect the flagellar transformation in ATCC10798 experiments, suggesting that the motor torque might be insufficient to cause flagellar transformation. To address this point, we quantified the motor properties using a tethered-cell assay (Fig. 3a, see Material and Methods). We recorded the rotation for 10 seconds in each measurement. The switching frequency and CW bias (CW time/total time) were 1.15 ± 0.60 s^−1^ and 0.61 ± 0.21, respectively, in ATCC10798; and 1.52 ± 0.84 s^−1^ and 0.48 ± 0.27 in the W3110 (Fig. 3b and c). The rotation rates of ATCC10798 and W3110 were 7.3 ± 1.4 Hz and 7.0 ± 1.3 Hz, respectively (Fig. 3d). We could not detect any difference in the motor speed between two strains (*P* = 0.1864 > 0.05 by *t*-test), suggesting that the defective flagellar polymorphism of ATCC10798 is not caused by its motor properties.

### Single-point mutation FliC(N87K) is essential for forward-backward movement

A bistable protofilament model explains the polymorphic flagellar transition. The flagellar filament is composed of 11 protofilaments, each of which assumes either a left- or right type, and this mixture of two types of protofilament produces several filament shapes, such as a normal, semi-coiled, and curly [34-36]. Additionally, it is known that some point mutations can lead to formation of these left- and right type protofilaments [37-39]. Therefore, we compared the *fliC* between ATCC10798 and W3110, and found that residue 87 of FliC in ATCC10798 was changed from asparagine to lysine (Fig. 4a). The effect of amino acid substitutions on polymorphic flagellar transformation is well studied, but the effect of this substitution on flagellar formation has never been investigated, to our knowledge.

To check whether this substitution was truly responsible for the transformation from a left-handed to right-handed flagellar filament, we replaced the *fliC* gene of ATCC10798 with a wild type one, SHU102 [ATCC10798(*fliC*(N87K)::*fliC*)] (see Material and Methods). We first examined the flagellar morphology using TEM (Fig. 4b). The pitch and helical radius of SHU102 flagella were 2.5 ± 0.2 μm (Fig. 4c *top*) and 0.20 ± 0.04 μm (Fig. 4c *bottom*), respectively, which corresponded to the normal flagellar type, as observed in W3110 (Fig. 1f and g *bottom*). We next investigated the swimming motility of SHU102. SHU102 formed a swarm ring on the semi-solid agar plate; and its diameter was similar to that observed in W3110 (Fig. 4d). Additionally, SHU102 cells displayed run-and-tumble strategy in its chemotactic behavior (Supplementary Videos 8-9 and Fig. 4e). Furthermore, we examined the effect of this substitution on chemotactic response, using a capillary (tip) assay, and found that FliC(N87K) substitution did not influence on it (Supplementary Video 10 and Supplementary Fig. 6). These results were confirmed independently by the experiment with Δ*fliC* cells expressing FliC(N87K) (Supplementary Result 1).

We also examined the rotation rate and morphology of the flagellar filaments using TIRFM (Supplementary Fig. 7) and found that SHU102 cells had the left-handed flagellar filament. The flagellar helicity frequently underwent switching into a right-handed form, depending on motor switching, which has never been observed in ATCC10798 cells (Supplementary Video 11). These results suggest that FliC(N87K) caused the structure of filaments to be fixed in a right-handed helicity.

### W3110 cells can escape from stuck on agarose surface through 180°-reverse movements

Although ATCC10798 cells show chemotaxis in a liquid environment (Supplementary Fig. 6), they were not able to swim on semi-solid agar (Fig. 1). To examine the reason, we checked swimming motility using agarose. As with the agar experiment, ATCC10798 cells could not form a swarm ring on a 0.2 % agarose plate, but W3110 cells could do so (Fig. 5a). Phase-contrast microscopy revealed that some W3110 cells were dispersed thinly to all areas (Fig. 5b (i) and (ii)), whereas ATCC10798 cells were more densely existed (Fig. 5b (iii)). To check this difference in detail, we observed the swimming motility of fresh cells using a 0.2 % semi-solid agarose pad (see Material and Methods). ATCC10798 cells were not able to swim once they stuck to the surface (Supplementary Video 12). On the other hand, W3110 cells frequently stuck to the surface but escaped via 180°-reversals, without reorientation of the cell body (Fig. 5d-f). Turner *et al* also observed the phenomenon using a fluorescent microscope: the flagellar bundle transformed from a normal to curly state, and the curly filaments formed a bundle that pushed the cell forward, in the opposite to the original direction of swimming. [40]. Additionally, we found that W3110 cells frequently reversed their swimming direction in the presence of 15 % (w/vol) Ficoll, as seen in constricted environments [41]; ATCC10798 cells were also able to swim with a forward and backward movement (Supplementary Video 13). Taken together, we conclude that flagellar polymorphism is one of the most crucial elements for migration in structured environments (see details in Discussion).

**Figure 5.**
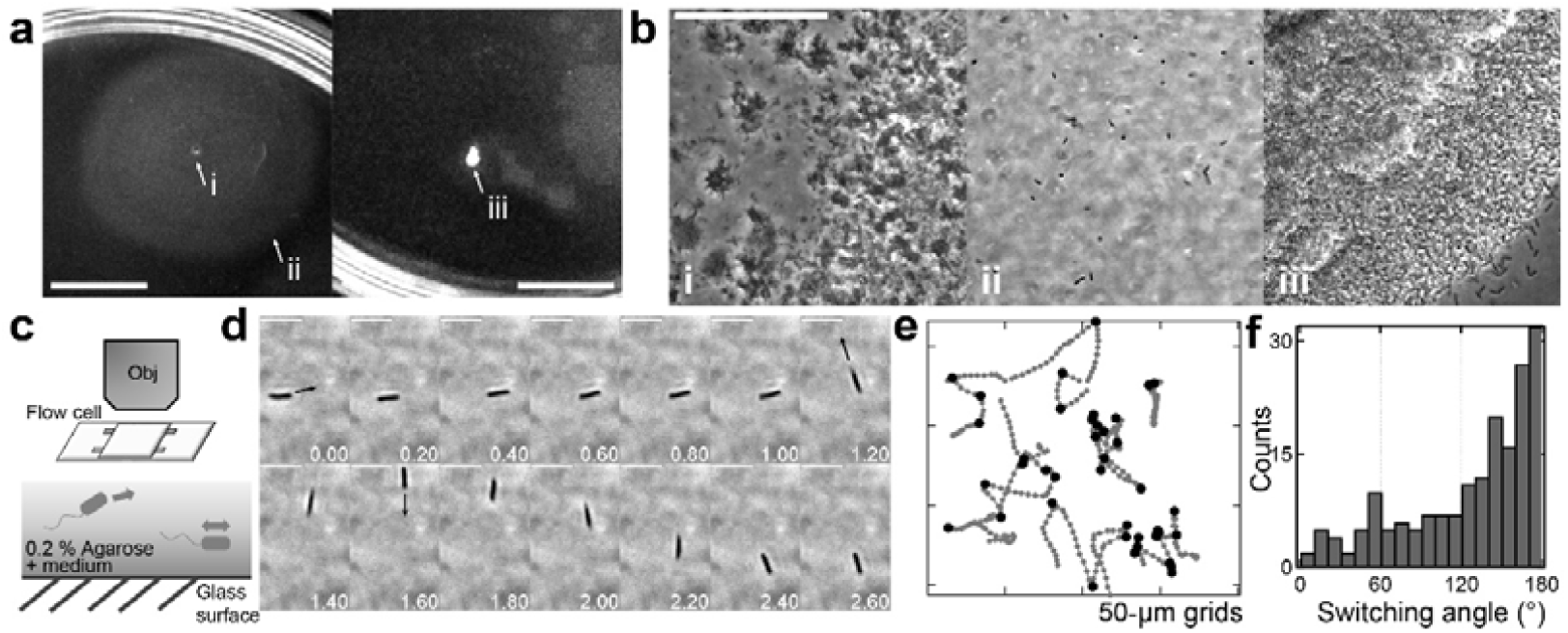
Swimming motility on 0.2 % agarose. (a and b) Motilities of W3110 cells (*left*) and ATCC10798 (*right*) on a 0.2 % (wt/vol) soft-agarose plate after incubation at 30 °C for 7 h. The magnified image at (i-iii) were shown in b. These experiments were conducted in the same plate. Scale bar, 1 cm (A) and 50 μm (b). (c) *Top*: The schematics to observe a swimming motility of W3110 cells in a 0.2 % soft agarose pad. *Bottom*: W3110 cells could swim in medium containing 0.2 % agarose. (d) Sequential images of migration in the 0.2 % agarose. Arrows indicate swimming directions after reversals, where the angle changes were approximately 180 degrees. Scale bar, 5 μm. (e) Typical examples of swimming trajectories with turn events. Black dots denote the time of reversals. Intervals, 20 ms. (f). Histogram of the switching angle (n = 183).

## Discussion

The FliC(N87K) substitution caused a flagellar transformation from the left-handed, normal flagellar filament into the right-handed, curly filament (Fig. 4). Flagellin monomer consists of four connected domains, D0-D3. The highly conserved D0 and D1 domains face inward into a filament core, while D2 and D3 domains protrude outside, against the central core (Supplementary Fig. 11a) [42]. The role of D2-D3 is to stabilize flagellar filaments. D0-D1 are mainly responsible for the L/R switching of protofilaments [42-44]. In the flagellar filaments of *S. typhimurium*, (and *E. coli*), amino acid substitution of D1 domains at A49, D108, D152, A415 (A417), A428 (A430), N434 (N436) and A450 (A452) causes a flagellar transformation from normal to curly state [37-39, 45]. These substitutions change the hydrogen bonding network for the L/R transition along 5-, 11- and 16-start filament interfaces. We checked the interactions between subunits using L-and R-type straight filaments of *S. typhimurium* [42]. Supplementary Fig. 11b highlighted their hydrogen bonding interactions. In the L-type filament, the E84 and E122 residues form multiple hydrogen bonds with the N439 residue at 5-start interface. In the R-type filament, hydrogen bonds are formed between E84 and T438 and between T130 and N439 at 5-start interface. These residues are conserved among different bacterial species (Supplementary Fig. 11c). Although the specific interaction of the N87 residue was not detected in both L- and R-type subunits, the N87K mutation is likely to cause the formation of new hydrogen bonds with the T438 residue at 5-start interface. This hydrogen bond might enhance the R-type interaction and cause the adoption of the right-handed helical form, as previously shown [38].

On a semi-solid agar plate, W3110 cells could form a ring, but ATCC10798 cells could not (Fig. 1). It is generally given that this ring formation is associated with chemotactic behavior, driven by a motor switching [28]. However, we infer that additional mechanisms are required for a ring formation, taking into consideration that ATCC10798 cells exhibit motor switching (Figs 2-3) and chemotactic behavior in liquid (Supplementary Fig. 6). Interestingly, W3110 cells exhibited 180°-reverse movements to escape from being stuck in a semi-solid agarose (Fig. 5), which is also observed in a peritrichous flagellated bacterium, *Bacillus subtilis* [46]. Turner *et al*. observed flagellar transformation-dependent 180° reversal movements using a fluorescent microscope (see Fig. 5 in [40]). This was only observed in structured environments [41, 46]. Considering that the flagellar morphology was stable, irrespective of motor switching (Supplementary Video 4), we propose a flagellar, polymorphism-dependent, migration mechanism in structured environments. Our proposal is supported by previous reports suggesting that a specific point mutation in FliC, which causes a lack of flagellar polymorphism, hinders the ability to swim on a semi-solid agar plate, but still allows movement in liquid media [37, 39].

We expect that the above model could also apply to other types of flagellated bacteria. Polar flagellated bacteria show flagellar polymorphic change from a normal to curly state in the single polar, flagellated species *Pseudomonas* spp, [47, 48] and from normal to coiled state for *Rhodobacter sphaeroides* [49]. A novel type of flagellar wrapping motion has recently been observed in the single polar, flagellated species *Shewanella putrefaciens* [11], multiple polar flagellated bacteria such as *Allivibrio fischeri, Burkholderia insecticola, and P. putida* [10, 50], and bipolar flagellated bacteria such as *Helicobacter suis* [51] and *Magnetospirillum magneticus* AMB-1 [52]. These bacteria reverse their direction of motion by the transition from CCW rotation of left-handed normal filaments into CW rotation of right-handed coiled filaments to escape from being trapped in structured environments. In *S. putrefaciens, flaB* is crucial, not only for flagellar polymorphism, but also the transition from regular swimming to wrapping motion. However, only FlaA cells are deficient in both motility and flagellar polymorphism [53]. In common with *S. putrefaciens*, polar-flagellated bacteria possess multiple flagellins for flagellar polymorphism and migration in structured environments [54-56]. These data support our idea that the ability of bacteria to swim in structured environments is driven by flagellar polymorphism. However, *Caulbacter cresentus* and *V. alginolyticus*, form a swarm ring on a semi-solid agar plate without flagellar polymorphism [55, 57]. In these bacteria, the hydrodynamic load causes the buckling of the straight hook, upon the motor switching from CW to CCW rotation [8, 9, 58]. This buckling mechanism could be equivalent to the flagellar polymorphism, as a means to perturb cell motile pattern. In fact, the poly-hook mutant of non-chemotactic cells forms a pseudo ring, driven by dynamic flagellar reorientation [59, 60].

What is the advantage of ATCC10798 cells possessing the curly filament, even though they have a risk of getting stuck in structured environments? Amino acid residues 90-97 in the N-terminal D1 domain of flagellin, conserved between β- and γ-proteobacteria, are essential to recognition by the innate immune receptors of host cells, known as “toll-like receptor 5” (TLR5) [61]. However, in α- and ε-proteobacteria, this amino acid sequence is altered, to escape from host recognition. These mutations abolish not only TLR5 recognition but also prevent motility on the semi-solid agar. Although the effect of the FliC(N87K) substitution on survival remains elusive, the alanine substitution FliC(L89A) does reduce TLR5 recognition by 50-60 % [62]. Because right-handed flagellar filaments could resist infection by bacteriophage χ [63, 64], we speculate that ATCC10798 cells have survived by the alteration of its flagellin sequence. An alternative theory is that cells without flagellar filaments are better at escaping detection by “predators”. However, antibacterial drugs also provide selective pressure for bacteria, in addition to phages and immune systems. To combat antibacterial drugs, it is known that some bacteria form biofilms. They attach to a surface, sticking to other bacteria via flagella and pili, then secrete extracellular polymeric substances, for homeostasis [65-67]. Considering that cells with curly filaments, unlike cells with normal flagellar filaments, easily adhere to one another [44] and aggregate in solution (Supplementary Fig. 6), we infer that ATCC10798 cells might have evolved curly filaments to support biofilm formation.

Taken together, we conclude that swarm ring formation corresponds to flagellar polymorphism and speculate that agar experiments fail to detect the motility of many cells. Our results complement recent, beautiful work on how microorganisms migrate in structured environments [18] and will lead to a discussion of how *E. coli* cells have adapted for survival through the evolution of flagellar transformation.

## Acknowledgements

The authors thank Prof. Keiichi Namba for use of his PyMOL model, Prof. Ritsu Kamiya in preparing the manuscript of the early version and Prof. Ikuro Kawagishi for fruitful discussions. This study was supported in part by the JSPS Funding Program for Next-Generation World-Leading Researchers Grant LR033 to T.N., by MEXT/JSPS KAKENHI Grants to T.N. (Nos. JP15H04364 and JP26103527) and to Y.S. (Nos. JP15K07034 and JP19H05404). Y.K was recipient of JSPS Fellowship for Japan Junior Scientists (15J12274) and the Uehara Memorial Foundation postdoctoral fellow.

## Author Contributions

Y.K. and Y.S. designed research; Y.K. performed research and collected data in R.B. and T.N. labs; T. I. collected a tethered cell data, M.Y., R.I, Y. V. M., and K.G helped for genetics and strain; T.N and Y.S developed the framework for analysis; Y.K., Y.V.M and Y.S. wrote the paper.

