## Supplementary Information for "Distinct chemotactic behavior in the original *Escherichia coli* K-12 depending on forward-and-backward swimming, not on run-tumble movements"

This file includes:

Supplementary Result 1

Supplementary Figures 1-11

Supplementary Table 1-3

Captions for Supplementary Videos

Supplementary References

**Supplementary Result 1**

**Effect of *fliC*(N87K) substitution on flagellar structure and swimming motility**

Using SHU102 [*fliC*(N87K)*::fliC*] cells, we demonstrated that the N87K substitution was truly responsible for the transformation from a left-handed to right-handed flagellar filament and for the change of swimming mode from a run-rumble to forward-backward swimming (Fig. 4).We furthermore examined the effect of the FliC(N87K) substitution by introducing the point mutation into wild-type *fliC* gene by PCR method (see Material and Methods). Using TEM (Supplementary Fig. 8a),we found that HCB1336 (Δ*fliC*) transformed with pSHU61 (pBR322-*fliC*(N87K)), called *fliC*(N87K) cells in the Supplementary information, formed approximately three filaments with a length of 4.5 ± 0.9 μm, and the flagellar pitch and helix radius were estimated to be 1.4 ± 0.1 μm and 0.14 ± 0.03 μm, respectively, which are similar to those of ATCC10798 cells (Supplementary Fig. 8b and c *top*). On the other hand, HCB1336 transformed with pYS10 (pBR322-*fliC*), called wild-type *fliC* cells in the Supplementary information, formed approximately three filaments with a length of 5.8 ± 1.6 μm around the cell body, and the flagellar pitch and helix radius were estimated to be 2.7 ± 0.3 μm and 0.22 ± 0.06 μm, respectively, which corresponded to the normal flagellar filament (Supplementary Fig. 8b and c *bottom*).

We next examined the effect of the FliC(N87K) substitution on the motility. *fliC*(N87K) cells did not form ring on a semi-solid agar plate, while wild-type *fliC* cells did (Supplementary Fig. 8d). In liquid mediums, wild-type *fliC* cells showed a linear motion with sporadic tumbling, while *fliC*(N87K) cells showed a wobble motion rather than a linear movement, like ATCC10798 cells (Supplementary Fig. 8e and Supplementary Video 8). Single-cell tracking revealed that the average swimming speed was 12.0 ± 4.2 μm s^-1^ in *fliC*(N87K) cells (Supplementary Fig. 8f *top*, n = 53) and 21.7 ± 4.5 μm s^-1^ in wild-type *fliC* cells (Supplementary Fig. 8f *bottom*, n = 53). For the turning angle, the peak in *fliC*(N87K) cells was approximately 140 degrees (Supplementary Fig. 8g *top*, n = 444), as shown in ATCC10798 (Fig. 1d *top*). On the other hand, the angle distribution in wild-type *fliC* cells was approximately 40° (Supplementary Fig. 8g *bottom*, n = 285), which was consistent with the value of standard *E. coli* strain [1]. Additionally, we found that both strains could sense the chemotactic gradients using a capillary (tip) assay; moreover, *fliC*(N87K) cells tend to aggregate like ATCC10798 cells (data not shown).

We then quantified the rotational rate and flagellar morphologies under TIRFM (Supplementary Video 11). The flagellar helicity was left-handed in wild-type *fliC* cells and right-handed in *fliC*(N87K) cells. The flagellar pitch and radius of *fliC*(N87K) cells were estimated to be 1.1 ± 0.1 μm and 0.14 ± 0.03 μm, respectively, which corresponded to the curly flagellar filament (Supplementary Fig. 9) [2]. The flagellar transformation was frequently detected in wild-type *fliC* cells, whereas *fliC*(N87K) cells did not show it like ATCC107908 cells. The parameters of structure and kinematics calculated with TIRFM were summarized in Supplementary Table 3. We also examined an effect of FliC(N87K) substitution on the motor properties using a tethered-cell assay and found no significant difference among strains (Supplementary Fig. 10).

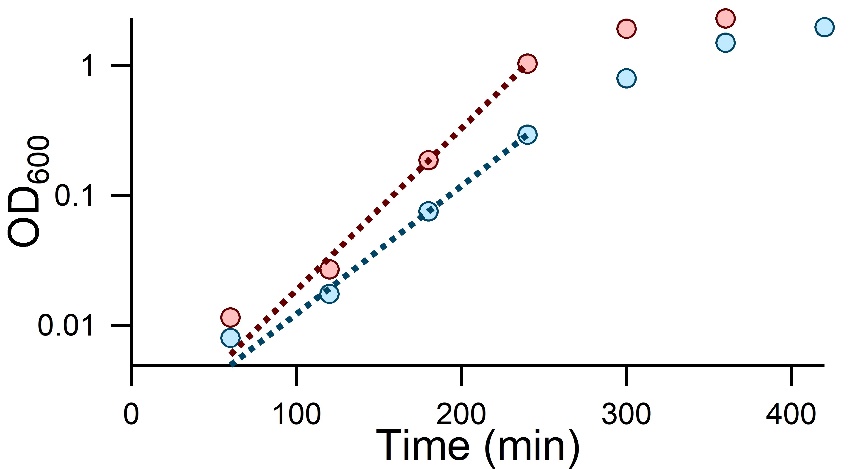

**Supplementary Figure 1 Growth curve in *Escherichia coli* K-12 ATCC10798 and W3110**

The single colony was scrutinized by tip, resuspended into 50-ml LB medium, and then cultured at 37°C with shaking. The absorbance of cells at 600 nm were periodically measured with 60-min intervals. During the exponential phase, we fitted the data as the following equation: f (t) = A×$2^{\frac{t}{tau}}$, where *A* the constant and *tau* the doubling time. The doubling time were estimated to be ~24 min in ATCC10798 (red) and 30 min in W3110 (blue). The experiments were performed two times.

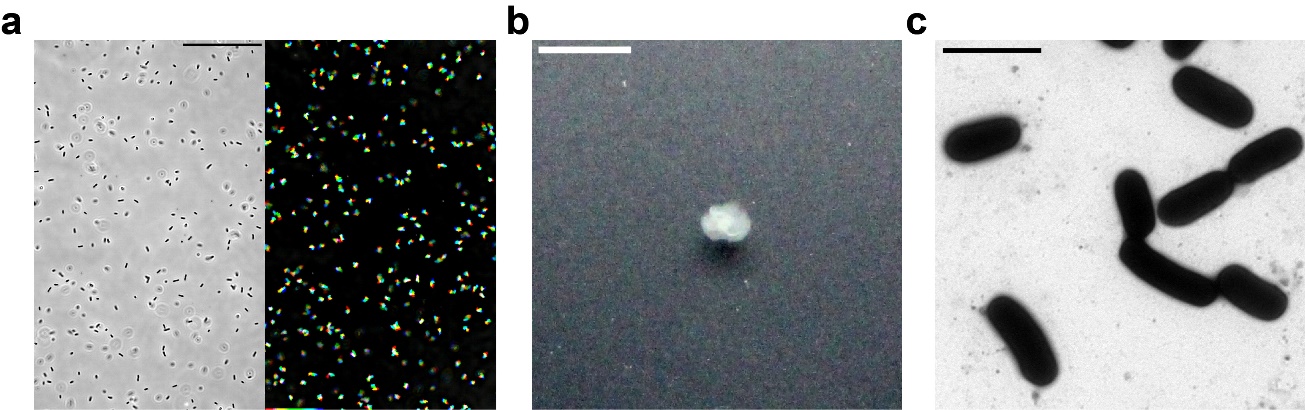

**Supplementary Figure 2 Characterization of swimming motility and structural parameters of SHU101**

(a) *Left*: Phase-contrast image of the non-flagellated mutant, SHU101. Scale bar, 50 μm. *Right*: The sequential phase-contrast images with 165-ms intervals were integrated for 5 s with the intermittent color code ‛red → yellow → green → cyan → blue.’ The processive linear movements could not be seen, indicating that the SHU101 is a non-motile strain. (b) Motility of SHU101 cells on a 0.25 % (wt/vol) soft-agar plate at 30ºC for 7 h. Scale bar, 0.5 cm. (c) Electron micrograph. Scale bar, 2 μm.

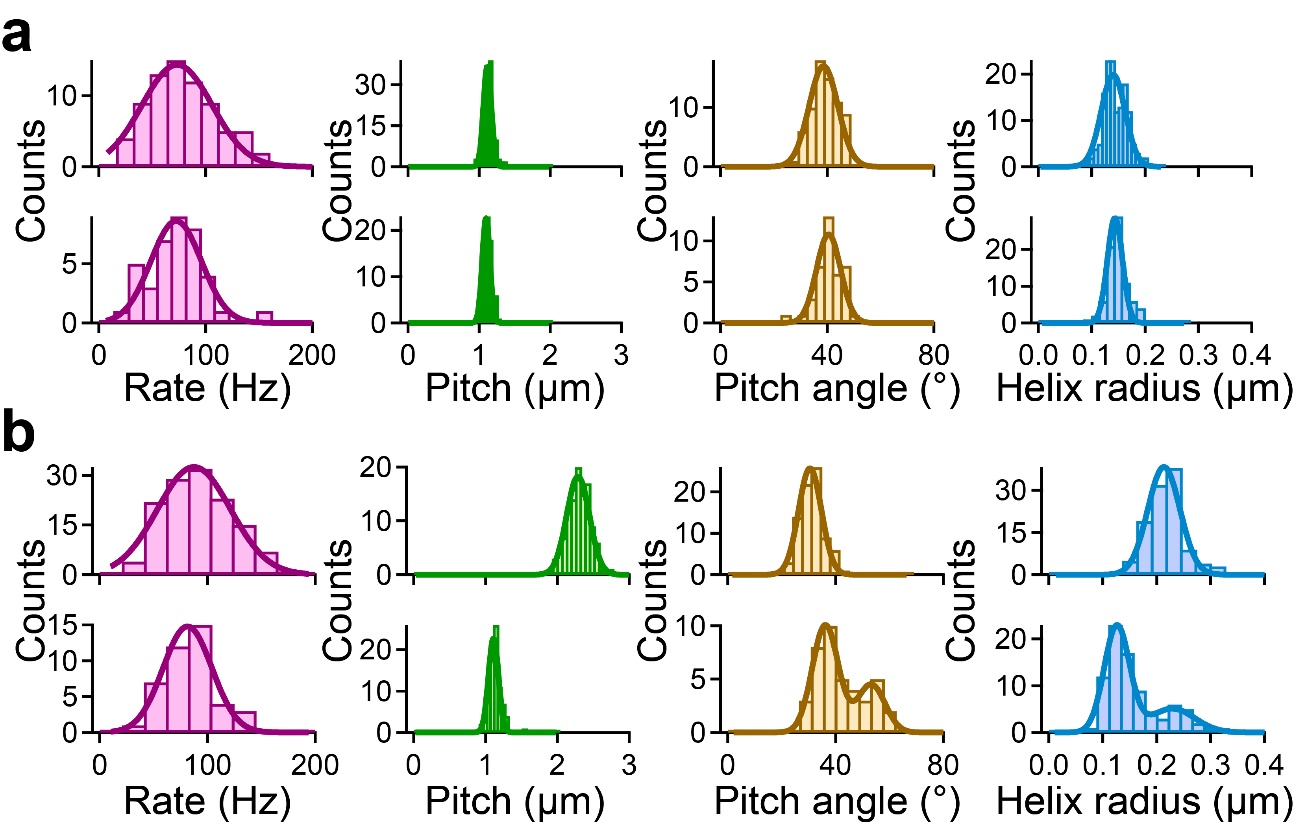

**Supplementary Figure 3 Quantification of rotational rate and structural parameters of flagella under total internal reflection fluorescence microscope (TIRFM).**

(a) Histograms of the rotation rate and structural parameters during CCW (*top*) and CW rotation (*bottom*) in ATCC10798 cells. Solid line represents the Gaussian fitting. The peaks and SDs of rotational rates were 73.3 ± 32.8 Hz in CCW direction (n = 74) and 72.4 ± 23.7 Hz in CW direction (n = 39). The flagellar pitches were 1.1 ± 0.1 μm in CCW (n =142) and 1.1 ± 0.1 μm in CW direction (n = 80). The pitch angles were 38.6 ± 5.4 degree in CCW (n = 70) and 40.5 ± 4.5 degree in CW (n = 39). The helix radii were 0.14 ± 0.02 μm in CCW (n = 142) and 0.14 ± 0.01 μm in CW (n = 80). (b) Histograms of the structure and rotation rate of flagella of W3110 during left-handed (*top*) and right-handed (*bottom*) state. The rotation rates were 87.6 ± 34.0 Hz under a left-handed state (n = 133) and 81.5 ± 22.7 Hz under a right-handed state (n = 42). The flagellar pitches were 2.3 ± 0.2 μm under a left-handed state (n = 112) and 1.1 ± 0.1 μm under a right-handed state (n = 84). The pitch angles were 30.5 ± 4.2 degree under a left-handed state (n = 81); 36.2 ± 4.8 and 53.2 ± 5.1 degree under a right-handed state (n = 40). The helix radii were 0.21 ± 0.03 μm under a left-handed state (n = 112); 0.13 ± 0.02 μm and 0.23 ± 0.04 μm under a right-handed state (n = 84).

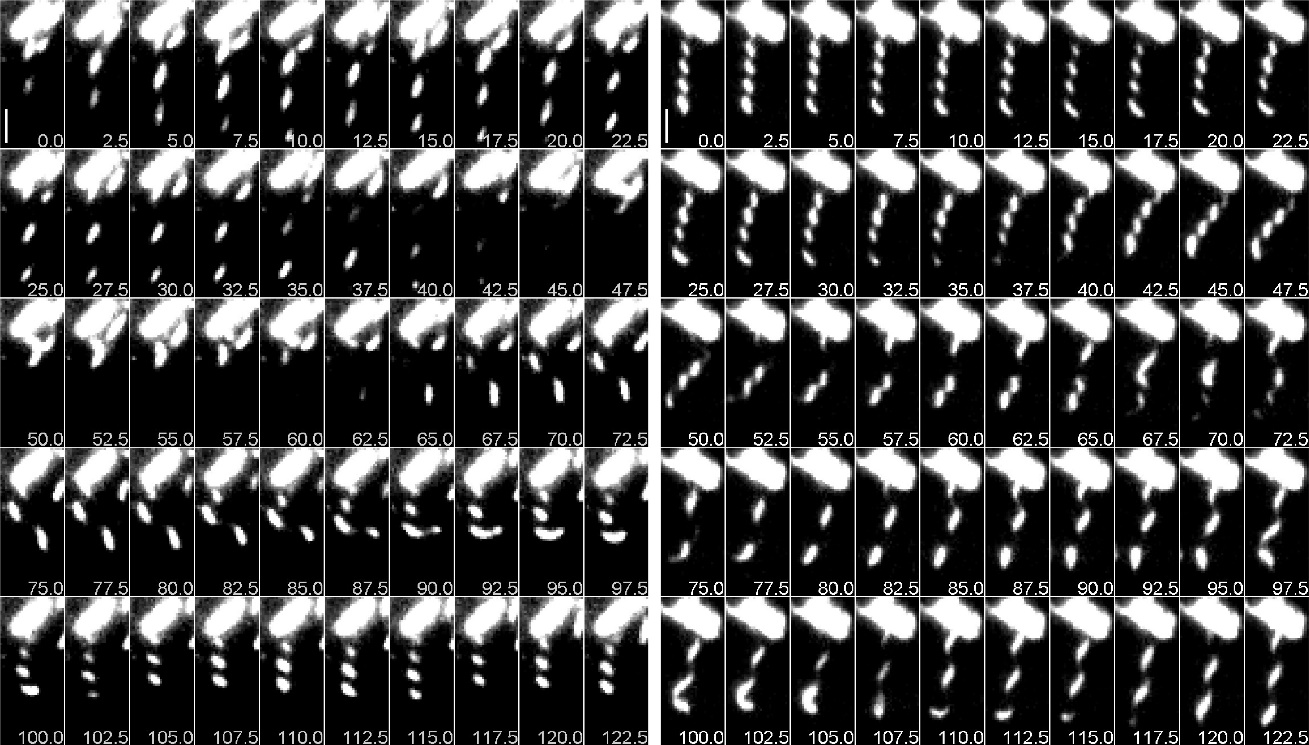

**Supplementary Figure 4 Real-time imaging of flagellar polymorphism in W3110**

Sequential images of the flagellar polymorphism at 2.5-ms intervals. *Left*: The orientation of flagellar filament(s) was from 1^st^ quadrant to 3^rd^ quadrant relative to the major axis of the filament, indicating that the helicity of filaments was left-handed. From 0 to 22.5 ms, the wave of flagella propagated in a direction away from the hook end toward the flagellar tip, indicating that the flagella rotated in CCW direction. The direction of rotation was switched at 30.0 ms, and then their helicity changed from the left-handed into right-handed within 50 ms. Taken together that the pitch angle of flagellar filaments was 58 degrees, the filament form was the semi-coiled. Scale bar, 2 μm. *Right*: The orientation of flagellar filament (s) was from 2^nd^ quadrant to 4^th^ quadrant relative to the major axis of the filament, indicating that the helicity of filaments was right-handed. We concluded that the flagellar form was the curly state with the fact of the 40°-pitch angle. From 0 to 17.5 ms, the flagellar filament did not move. At 20 ms, the flagellar filaments gradually moved, suggesting that the flagella started to rotate. The flagellar filaments dynamically twisted at 40 ms, and then the right-handed and left-handed flagellar were combined into a single filament at 67.5 and 105.0 ms, which was also seen in another bacterium. Finally, the curry filament(s) transformed into the normal state within 100 ms. Scale bar, 2 μm. Data from Supplementary Videos 6 and 7.

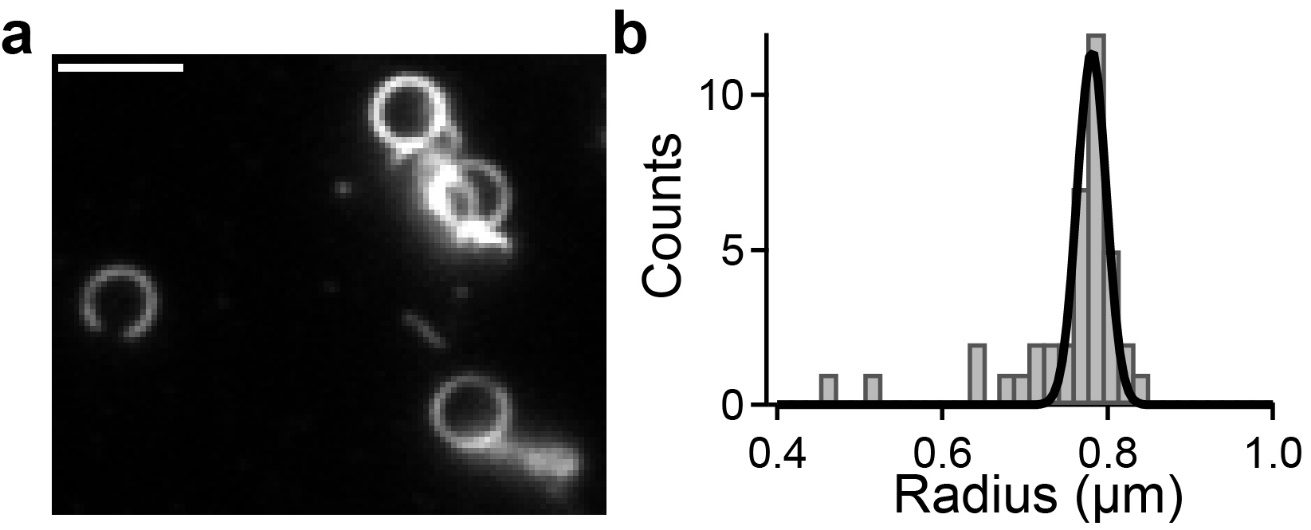

**Supplementary Figure 5 Quantification of flagellar radius at coiled state in W3110**

(a) The fluorescent micrograph of coiled-state flagellar filaments. Scale bar; 3 μm. (b) Histogram of the radius of coiled flagella. The solid line represents the Gaussian fitting, where the peak and SD are 0.78 ± 0.02 μm (n = 39).

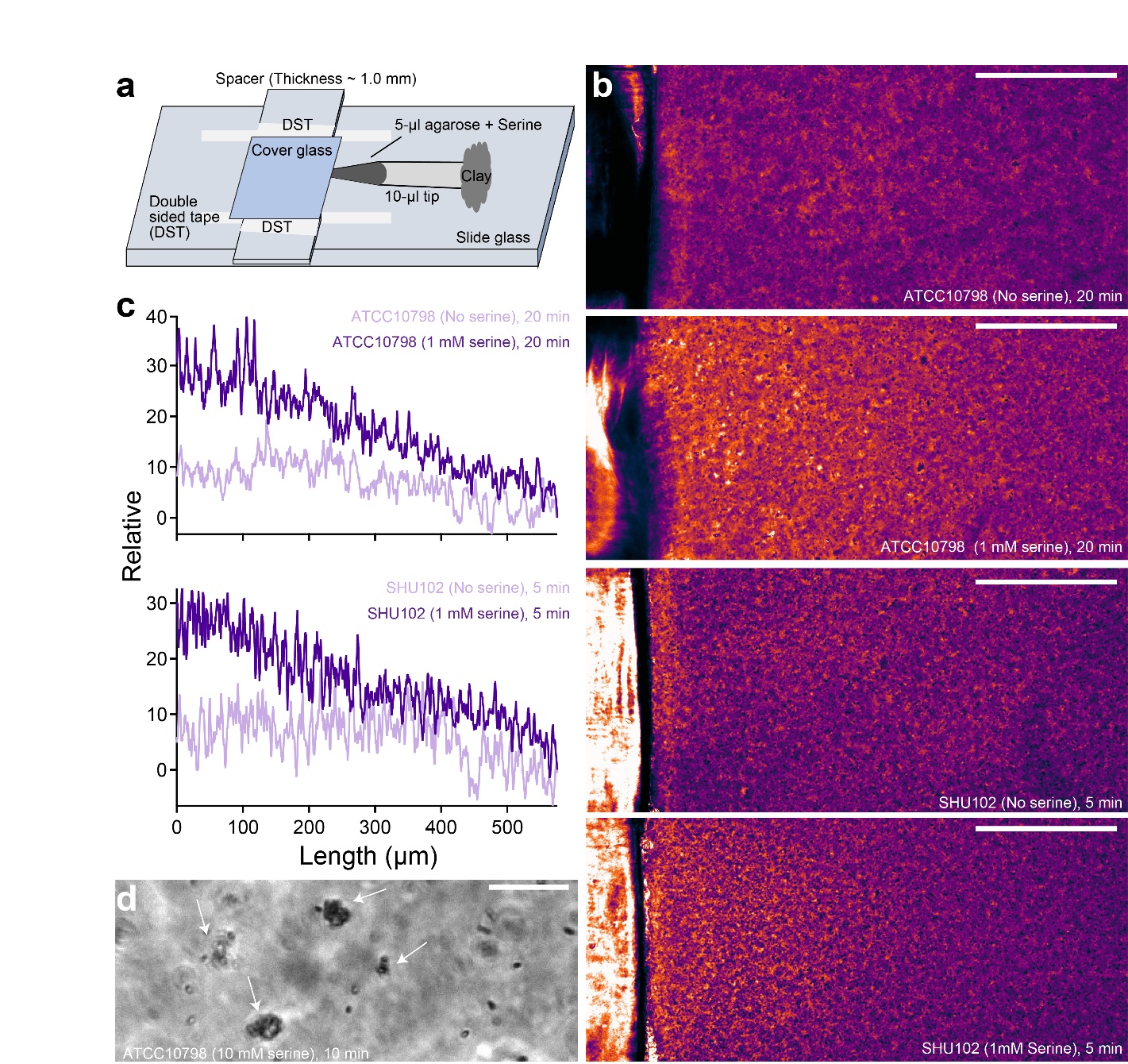

**Supplementary Figure 6 Chemotactic response of ATCC10798 [*fliC*(N87K)] and SHU102 [*fliC*(N87K)::*fliC*] cells**

(a) The schematic of tip (capillary) assay. The tip contains 5-μl buffer containing 1 % (wt/vol) agarose, and the other end was sealed with cray to avoid an effect of oxygen on a chemotactic response. (b) Pseudo-colours of phase-contrast images of ATCC10798 (*top*) and SHU102 cells (*bottom*). In the presence of 1 mM serine, both *E.coli* strains exhibit a chemotactic response and gather near a tip (orange color). Scale bar, 200 μm. (c) Intensity profiles of b. (d) A phase-contrast image of ATCC10798 cells in the presence of 10 mM serine. In the presence of serine, ATCC10798 cells tends to gather and aggregate each other, which was not detected in SHU102 cells. Scale bar, 20 μm.

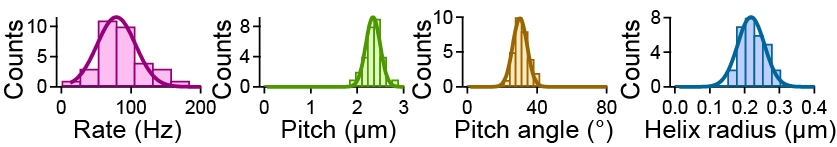

**Supplementary Figure 7 Quantification of rotational rate and structural parameters of flagella of SHU102 under TIRFM**

Histograms of the structure and rotation rate of flagella of SHU102. The solid line represents the Gaussian fitting. Peaks and SDs of the rotation rate were 78.9 ± 27.5 Hz (n = 33). The flagellar pitch was 2.3 ± 0.2 μm (n = 30). The pitch angles were 30.2 ± 4.1 degree (n = 30). The helix radius was 0.22 ± 0.04 μm (n = 30).

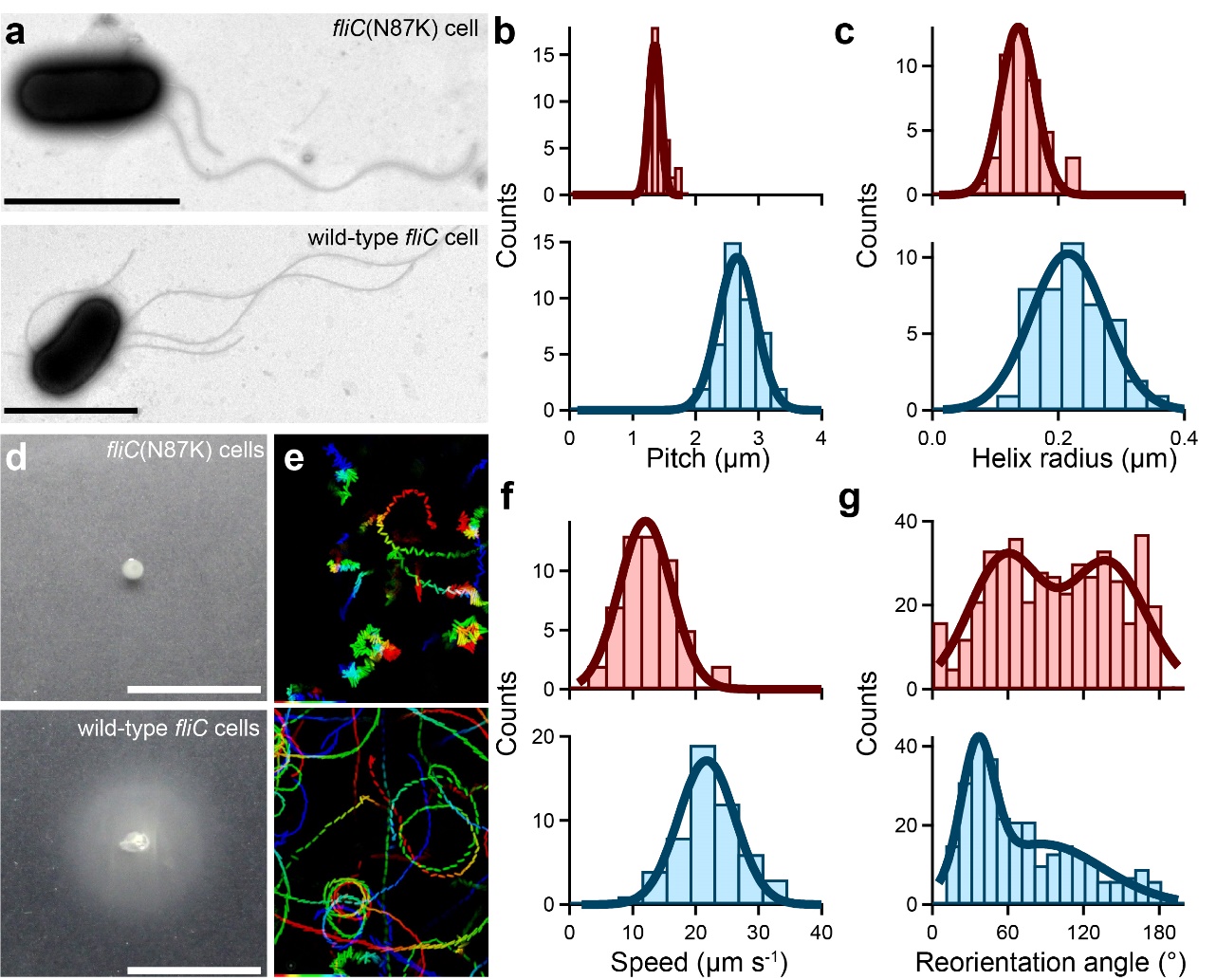

**Supplementary Figure 8 Comparison of the flagellar structure and swimming motility between wild-type *fliC* and *fliC*(N87K) cells**

(a) Electron micrograph of *E. coli* cells. Scale bars, 2 μm. (b) Histogram of the pitch. The solid lines represent the Gaussian fitting, where the peaks and SDs are 1.4 ± 0.1 μm in *fliC*(N87K) cells (*top*, n = 46) and 2.7 ± 0.3 μm in *fliC* cells (*bottom*, n = 42). (c) Histogram of the helix radius. The peaks and SDs are 0.14 ± 0.03 μm in *fliC*(N87K) cells (*top*, n = 46) and 0.22 ± 0.06 μm in *fliC* cells (*bottom*, n = 44). (d) Motilities on the 0.25 % (wt/vol) soft-agar plates at 30ºC for 7 h. Scale bar, 1 cm. (e) Swimming traces at 150-ms intervals for 15 s. The intermittent color code indicated the time course from red to blue. Area, 68.6 μm × 85.9 μm. (f) Histograms of the swimming speed. The peaks and SDs are 12.0 ± 4.2 μm s^-1^ in *fliC*(N87K) cells (*top*, n = 53) and 21.7 ± 4.5 μm s^-1^ in *fliC* cells (*bottom*, n = 53). (g) Histograms of reorientation angles. The peaks and SDs are 58 ± 29 degree and 139 ± 30 degree in *fliC*(N87K) cells (*top*, n = 444); 36 ± 13 degree in *fliC* cells (*bottom*, n = 285).

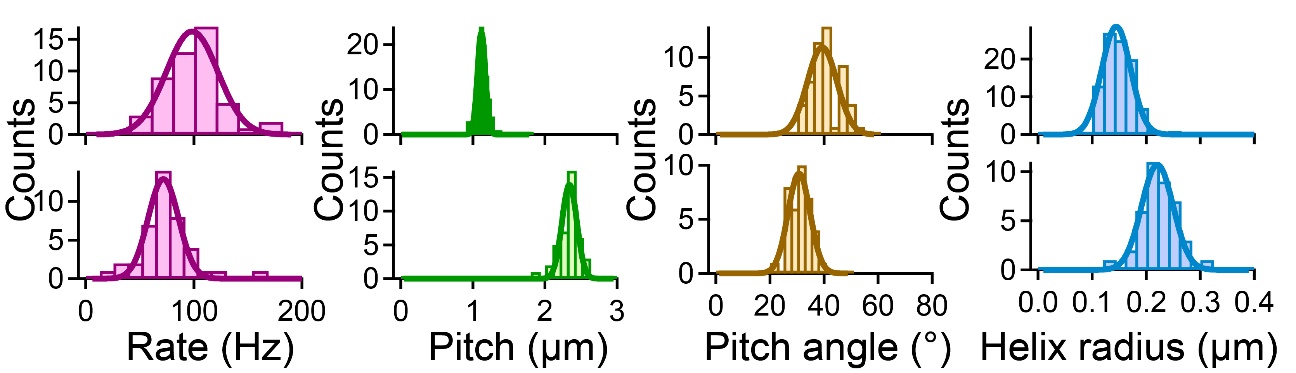

**Supplementary Figure 9 Quantification of rotational rate and structural parameters of wild-type FliC and FliC N87K under TIRFM**

Histograms of the structure and rotation rate of flagella of *E. coli* Δ*fliC* strain with the plasmid encoding wild-type *fliC* and *fliC*(N87K). Solid line represents the Gaussian fitting. Peaks and SDs of the rotation rate were 98.4 ± 23.9 Hz in *fliC*(N87K) cells (*top*, n = 50) and 71.8 ± 14.1 Hz in wild-type *fliC* cells (*bottom*, n = 41). The flagellar pitches were 1.1 ± 0.1 μm in *fliC*(N87K) cells (top, n = 94) and 2.3 ± 0.1μm in wild-type *fliC* cells (*bottom*, n = 40). The pitch angles were 39.3 ± 5.5 degree in *fliC*(N87K) cells (*top*, n = 52) and 30.8 ± 4.1 degree in wild-type *fliC* cells (*bottom*, n = 37). The helix radii were 0.14 ± 0.03 μm in *fliC*(N87K) cells (*top*, n = 94) and 0.22 ± 0.03 μm in wild-type *fliC* cells (*bottom*, n = 40).

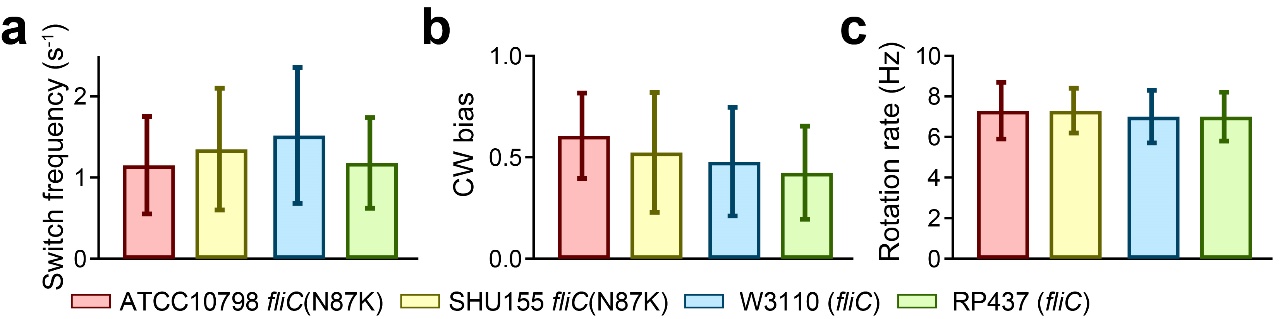

**Supplementary Figure 10 Quantification of motor properties by tethered-cell assay**

(a) Switching frequency between *E. coli* strains for 10 sec. The average and SD were 1.15 ± 0.60 s^-1^ in ATCC10798 (n = 99), 1.35 ± 0.75 s^-1^ in SHU155 (n = 67), 1.52 ± 0.84 s^-1^ in W3110 (n = 53), and 1.18 ± 0.56 s^-1^ in RP437 (n = 49). (b) CW bias (Time_CW_/Time_Total_). The average and SD were 0.61 ± 0.21 in ATCC10798 (n = 99) and 0.52 ± 0.30 in SHU155 (n = 67), 0.48 ± 0.27 in W3110 (n = 53), and 0.42 ± 0.23 in RP437 (n = 49). (c) Rotation rates. The average and SD were 7.3 ± 1.4 Hz in ATCC10798 (n = 99), 7.3 ± 1.1 Hz in SHU155 (n = 67), 7.0 ± 1.3 Hz in W3110 (n = 53), and 7.0 ± 1.2 Hz in RP437 (n = 49). ATCC10798 and W3110 data from Figure 3.

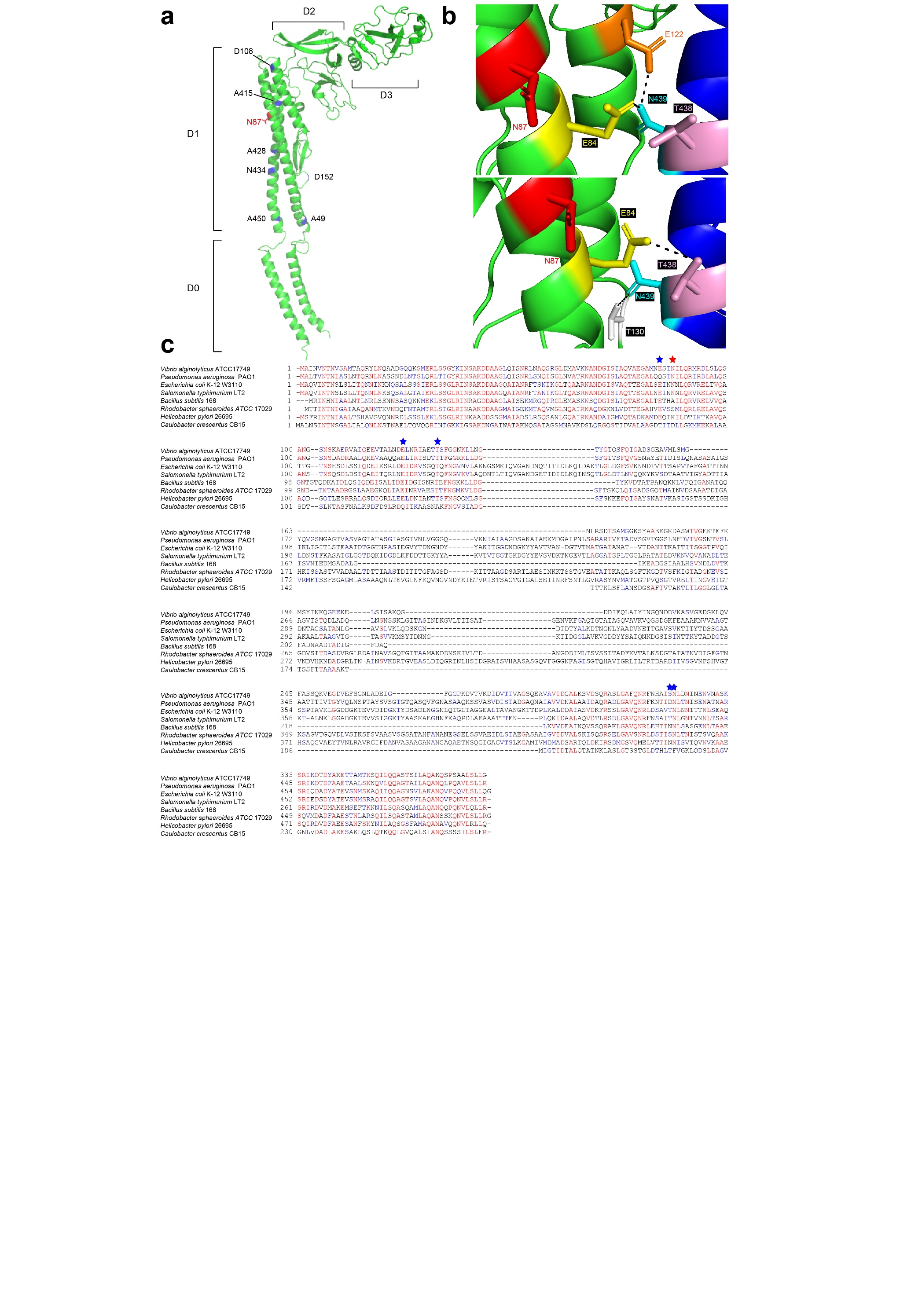

**Supplementary Figure 11 Structure and sequence of flagellin**

(a) Subunit structure of L-type flagellar filament. The important residues for a curly filament are shown. (b) L-type (*top*) and R-type (*bottom*) straight filaments of *S. typhimurium* are shown [3]. Hydrogen bonds interaction (black) between S0 (green) and S5 subunits (blue). E84, N87, E122, T130 of S0 subunit are colored in yellow, red, orange, and white, respectively; T438 and N439 of S5 subunit are colored in cyan and pink, respectively. (c) Alignments of the ﬂagellin amino acid sequence from typical bacteria. Red star represents N87 residue; blue stars indicate E84, E122, T130, T438, N439 residues.

**Supplementary Table 1. Strains and plasmids**

| Strain/plasmid | Relevant Phenotype/ Genotype | Note |
| --- | --- | --- |
| *Strains* |  |  |
| W3110 | Wild type |  |
| ATCC10798 | Wild type (*fliC* N87K) | This study |
| RP437 | Wild type for chemotaxis | [4] |
| HCB1336 | CM735 Δ*fliC* |  |
| SHU101 | ATCC10798 *fliC*(N87K)::*tetRA* | This study |
| SHU102 | ATCC10798 *fliC*(N87K)::*fliC* | This study |
| SHU155 | W3110 *fliC*::*fliC*(N87K) | This study |
| *Plasmids* |  |  |
| pKD46 | Red system | [5] |
| pYS10 | FliC expression plasmid; pBR322 derivative | [6] |
| pSHU61 | FliC(N87K) expression plasmid; pBR322 derivative | This study |

**Supplementary Table 2. Primers**

| Name | 5’ > 3’ | Note |
| --- | --- | --- |
| FliC ATCC FW | GGAAACCCAATACGTAATCA | Forward primer for PCR and sequencing of *fliC* in ATCC10798 |
| FliC ATCC Rev | CAATTTGGCGTTGCCGTCAGTCTC | Reverse primer for PCR and sequencing of *fliC* in ATCC10798 |
| 1217_fliC(N87K)-f(QC) | CTGTCCGAAATCAACAAGAACTTACAGCGTGTG | Forward primer for mutagenesis on *fliC* |
| 1218_fliC(N87K)-r(QC) | CACACGCTGTAAGTTCTTGTTGATTTCGGACAG | Reverse primer for mutagenesis on *fliC* |
| 0196_fliC-tetRA-F | CAATATAGGATAACGAATCATGGCACAAGTCATTAATACCTTAAGACCCACTTTCACATTT | Forward primer for PCR of *tetRA* cassette |
| 0197_fliC-tetRA-R | ACCCTGCAGCAGAGACAGAACCTGCTGCGGTACCTGGTTACTAAGCACTTGTCTCCTG | Reverse primer for PCR of *tetRA* cassette |
| 1232_fliC-F | CAATATAGGATAACGAATCATGGCACAAGT | Forward primer for PCR of wild-type *fliC* cassette |
| 0199_fliC-R | TTAACCCTGCAGCAGAGACAGA | Reverse primer for PCR of wild-type *fliC* cassette and sequencing of *fliC* |
| 0210_tetRA-785-R | GGCAAGACTGGCATGATAAGGCC | Reverse primer for Colony PCR |
| 0211_tetRA-1090-F | GTGAAGTGGTTCGGTTGGTTAGGG | Forward primer for Colony PCR |
| 0219_fliC-(-175)-F | ATAGCGGGAATAAGGGGCAGA | Forward primer for Colony PCR and sequencing of *fliC* |
| 0220_fliC- (+250)-R | GGTGGCGGGGAAGCACGTTGC | Reverse primer for Colony PCR and sequencing of *fliC* |
| 0198__fliC-F | ATGGCACAAGTCATTAATACC | Forward primer for sequencing of *fliC* |

**Supplementary Table 3**

Structural parameters and kinematics of the cell body and flagella in *Escherichia coli*. Values were directly measured by either a transmission electron microscopy or optical microscopy.

| Cells | Body length  (μm) | Body width (μm) | Flagellar length  (μm) | Flagellar helix radius  (μm) | Flagellar  number | Flagellar pitch  (μm) | Swimming speed  (μm s^-1^) | Rotation  rate  (Hz) |
| --- | --- | --- | --- | --- | --- | --- | --- | --- |
| ATCC10798 | 1.9±0.3  (62) | 0.70±0.07  (62) | 4.7±1.1  (62) | 0.14±0.03  (59) | 1.8±0.6  (61) | 1.3±0.2  (59) | 13.2±4.4  (70) | 73.3±32.8  (74) |
| W3110 | 2.1±0.5  (50) | 0.77±0.13  (50) | 7.3±1.9  (48) | 0.22±0.05  (42) | 6.5±2.3  (39) | 3.0±0.2  (41) | 32.5±6.6  (50) | 87.6±34.0  (133) |
| pYS10/HCB1336 | 2.1±0.4  (57) | 0.83±0.17  (57) | 5.8±1.6  (57) | 0.22±0.06 (44) | 3.2±1.3  (57) | 2.7±0.3  (42) | 21.7±4.5  (53) | 71.8±14.1  (41) |
| pSHU61/HCB1336 | 1.8±0.2  (46) | 0.73±0.12  (46) | 4.5±0.9  (44) | 0.14±0.03  (46) | 3.1±1.3  (46) | 1.4±0.1 (46) | 12.0±4.2  (53) | 98.4±23.9  (50) |
| SHU101 | 1.9±0.3  (21) | 0.74±0.06  (21) | n.d. | n.d. | 0  (21) | n.d. | n.d. | n.d. |
| SHU102 | 2.1±0.3  (32) | 0.95±0.20  (32) | 6.9±1.5  (29) | 0.20±0.04  (27) | 5.3±2.4  (31) | 2.5±0.2  (27) | 26.3±6.0  (45) | 78.9±27.5  (33) |
|  |  |  | EM | |  |  | OM | |

EM, estimation from data taken by electron microscopy; OM, estimation from sequential images by the optical microscopy.

**Captions for Supplementary Videos**

Supplementary Video 1

Swimming motility of *E. coli* K-12 ATCC10798 (*left*) and *E. coli* K-12 W3110 (*right*) observed under a phase-contrast microscope. ATCC10798 cells show forward-and backward movements, whereas W3110 cells show run and tumble movements. Scale bar, 20 μm.

Supplementary Video 2

Swimming motility of ATCC10798 (*left*) and W3110 (*right*) under a conventional fluorescent microscope. Scale bar, 20 μm.

Supplementary Video 3

High-speed imaging of flagellar rotation of ATCC10798 (*left*) and W3110 (*right*) using an EMCCD camera under a fluorescent microscope. Scale bar, 5 μm

Supplementary Video 4

Flagellar rotation of ATCC10798 under TIRFM. Scale bar, 4 μm

Supplementary Video 5

Flagellar rotation of W3110 under TIRFM. Scale bar, 5 μm

Supplementary Video 6

Real-time imaging of a polymorphic flagellar change under TIRFM. A rotational direction was changed at 0.11 sec; subsequently, a normal left-handed filament was transformed into the right-handed semi-coilded filament. Scale bar, 2 μm.

Supplementary Video 7

Real-time imaging of a polymorphic flagellar change under TIRFM. A rotational direction was changed at 0.17 sec; subsequently, a right-handed curly filament was transformed into the normal left-handed filament. Scale bar, 2 μm.

Supplementary Video 8

Swimming motility wild-type pYS10/HCB1336 (*left*), pSHU61/HCB1336 (*middle*)*,* and SHU102 cells (*right*) under a phase-contrast microscope. Scale bar, 40 μm.

Supplementary Video 9

Swimming motility of SHU102 cells under a conventional fluorescent microscope. Scale bar, 20 μm.

Supplementary Video 10

Chemotactic response of ATCC10798 (*top*) and SHU102 cells (*bottom*) under a phase-contrast microscope. For a first 10 sec, cells swim in the absence of serine. For a last 10 sec, both cells swim in the presence 1 mM serine. Scale bar, 200 μm.

Supplementary Video 11

Flagellar roration of pYS10/HCB1336 (*left*), pSHU61/HCB1336 (*middle*)*,* and SHU102 cells (*right*) under TIRFM. Scale bar, 2 μm.

Supplementary Video 12

Swimming motility of ATCC10798 (*left*) and W3110 (*right*) on a 0.2 % agarose pad. ATCC10798 cells cannot migrate after stuck, whereas W3110 cells keep moving with 180°-reversals to escape from a stuck. Scale bar, 50 μm.

Supplementary Video 13

Swimming motility of ATCC10798 (*left*) and W3110 (*right*) in the presence of 15 % ficoll. Scale bar, 20 μm.
